# Early occurrence of *Teratoramularia rumicicola* in Japan: re-identification of historic strain GR1 and its pathogenic effects on *Rumex* species

**DOI:** 10.64898/2025.12.22.696101

**Authors:** Masataka Izumi, Toyozo Sato

**Affiliations:** Graduate School of Agriculture, Kyoto University, Kyoto, Japan; Niigata Agro-Food University, Niigata, Japan

**Keywords:** *Teratoramularia rumicicola*, *Rumex japonicus*, re-identification, natural host, historical record

## Abstract

*Teratoramularia rumicicola* is a plant pathogenic fungus infecting *Rumex* species. Although it was described as a new species in 2016, its fundamental characteristics, including historical occurrence, geographic distribution, and host range, remain poorly resolved. In this study, we re-identified strain GR1, which was originally isolated from a diseased *Rumex* plant in the Bonin (Ogasawara) Islands in 1996 and deposited in a public bioresource center, together with the plant voucher specimen, and evaluated its host range. Colony and conidial morphology and a phylogenetic analysis based on concatenated internal transcribed spacer (ITS) and large subunit (LSU) rDNA sequences identified GR1 as *T. rumicicola*. Morphological examination of the host plant voucher specimen confirmed *Rumex japonicus* as the natural host. Pathogenicity assays demonstrated that GR1 infected *R. japonicus*, reproduced the original disease symptoms, and was successfully re-isolated from diseased tissues, consistent with Koch’s postulates. In addition, GR1 caused significant disease and shoot biomass reduction in *R. japonicus*, *R. crispus* and *R. obtusifolius*. The identification of GR1 reveals the presence of *T. rumicicola* in Japan approximately two decades prior to its formal description, representing the earliest known record of this species. Furthermore, this finding extends its known distribution from temperate regions to the subtropical Bonin Islands, a well-preserved and isolated ecosystem. This study documents for the first time *R. japonicus* as a natural host of *T. rumicicola*.

## Introduction

The ramularioid fungi, historically accommodated in *Ramularia*, represents a morphologically heterogeneous group of hyphomycetes with ramularia -like conidiogenesis and both economic and ecological importance (Videira et al. 2016). Many of these species are economically significant foliar pathogens of cereals, forage grasses, and other crops, causing leaf spot and blight diseases that reduce yield and quality, with *Ramularia collo-cygni* B. Sutton & J.M. Waller on barley as a notorious example (McGrann et al. 2016). In addition to their importance as crop pathogens, several *Ramularia*-like fungi have also attracted attention as potential agents for the biological control of weeds (Bond et al. 2007), highlighting their dual relevance to both plant pathology and weed management. Given this agricultural significance, accurate delimitation of these fungi is essential. In this context, molecular phylogenetic analyses revealed the ramularioid fungi to be polyphyletic, leading to the erection of several distinct genera, including *Teratoramularia* (Videira et al. 2016).

*Teratoramularia* currently comprises several species, including *Teratoramularia rumicicola* Videira, H.D. Shin & Crous, *Teratoramularia rumicis* Kushwaha, S.K. Verma, Sanj. Yadav & Raghv. Sing, *Teratoramularia kirschneriana* Videira & Crous, *Teratoramularia infinita* Videira & Crous, and *Teratoramularia persicariae* Videira, H.D. Shin & Crous. Among them, *T. rumicicola* is recognized as a foliar pathogen of problematic *Rumex* weeds and has been highlighted as a promising candidate for microbial bioherbicide development in pasture systems (Videira et al. 2016; Choi et al. 2025; Izumi and Sato 2025) . However, as a species described only recently, its fundamental characteristics—particularly historical occurrence, geographic distribution, and host range —remain poorly understood. To date, *T. rumicicola* has been reported from *Rumex crispus* L. in Korea and mainland Japan (Videira et al. 2016; Izumi and Sato 2025) and from *Rumex obtusifolius* L. in Korea (Choi et al. 2025). Artificial inoculations with Japanese isolates further demonstrated the susceptibility of *Rumex japonicus* Houtt., although this species has not yet been documented as a natural host (Izumi and Sato 2025). Overall, current information on the host range and ecology of *T. rumicicola* remains limited. Clarifying such fundamental traits is essential not only for refining the taxonomy of this newly described species, but also for understanding its biogeography and assessing its potential applications in weed management.

In this study, we focus on strain GR1, collected in 1996 from a diseased *Rumex* plant on Hahajima, the Bonin Islands, by T. Sato (a co-author). It was initially listed as *Ramularia pratensis* Sacc. in a non-peer-reviewed report (Sato et al. 2010) and deposited in the NARO Genebank as MAFF 238946. An internal transcribed spacer (ITS) sequence later suggested affinity with *T. rumicicola* or *T. rumicis*, but the isolate itself has never been formally reassessed. Moreover, although recorded as originating from *R. japonicus*, no morphological evidence was provided for the host plant, leaving the authenticity of this host record uncertain. Thus, both the fungal identity and its host association remain unresolved, underscoring the need for formal reassessment.

The objectives of this study were to re-examine strain GR1 using ITS and LSU sequence analyses and morphological observations, to reassess the host voucher morphologically, and to evaluate the pathogenicity of GR1 against *Rumex* species. These investigations aim to clarify the taxonomic placement of GR1 and its host association, thereby providing new insights into the historical presence and ecological breadth of *T. rumicicola* in Japan.

## Materials and Methods

### Fungal Strain GR1

Strain GR1 was originally isolated in June 1996 by coauthor T. Sato from a naturally infected *Rumex* leaf exhibiting white powdery mildew-like symptoms in Hahajima in the Bonin Islands, Japan. At the time of isolation, the fungus was recorded as *R. pratensis* and deposited in the NARO Genebank under accession MAFF 238946. For the present study, GR1 was revived from the culture collection and used for morphological, molecular, phylogenetic, and pathogenicity analyses.

### Morphological Observations of Strain GR1

Morphological examination of strain GR1 followed the procedures described previously with minor modifications (Izumi and Sato 2025). Briefly, GR1 was grown on potato dextrose agar (PDA; Shimadzu Corporation, Kyoto, Japan) at 25 °C in the dark. Colony characteristics were recorded after 33 days of incubation, whereas conidia for microscopic observation were obtained from 24 -day-old colonies. Conidial morphology, including length and septation, was examined under a light microscope (BH-2, Olympus, Japan). Morphological characteristics were evaluated by reference to published descriptions of *T. rumicicola* and *T. rumicis* (Videira et al. 2016; Verma et al. 2021).

### Molecular and Phylogenetic Analyses of Strain GR1

Molecular and phylogenetic analyses followed the methods described previously (Izumi and Sato 2025). Briefly, genomic DNA of GR1 was extracted from mycelium grown on PDA at 25 °C in the dark, and ITS and large subunit (LSU) regions were amplified using the primer pairs ITS1/ITS4 (White et al. 1990) and LR0R/LR7 (Vilgalys and Hester 1990), respectively. Amplicons were purified and sequenced bidirectionally. Consensus sequences were assembled using CodonCode Aligner v12.0.1 and deposited in GenBank under accessions PX601562 (ITS) and PX599000 (LSU). The resulting sequences were analyzed using BLASTn on the NCBI website to identify their closest taxonomic affiliations.

ITS and LSU sequences of strain GR1 were concatenated and aligned with representative *Teratoramularia* species using MUSCLE in MEGA v11 (Tamura et al. 2021). Ambiguously aligned regions were removed, yielding a final alignment of 1,191 bp. The maximum-likelihood phylogeny was reconstructed in RAxML v8.2.4 (Stamatakis 2014) using the GTR+GAMMA substitution model with 1,000 bootstrap replicates. The tree was rooted with *Staninwardia suttonii* Crous & Summerell CPC 13055 and displayed in iTOL (Letunic and Bork 2024). Sequences retrieved from GenBank are listed in Table S1.

### Host Plant Voucher Specimen

The host plant voucher specimen corresponding to strain GR1 was collected in June 1996 on Hahajima in the Bonin Islands, Japan, by coauthor T. Sato. At the time of collection, the plant was recorded as *R. japonicus* without any accompanying morphological description. The specimen is currently preserved in the herbarium of the Institute for Agro-Environmental Sciences, now a part of NARO, under voucher

ID NIAES-H 20884. For this study, the voucher was retrieved from the herbarium collection and used for morphological observations.

### Morphological Examination of the Voucher

Morphological observations of the host voucher specimen were conducted to confirm the host identity. Diagnostic characters examined included the leaf base and the inner tepal, which are regarded as key traits for distinguishing *Rumex* species (Ohashi et al. 2017). In particular, the margin of the inner tepal is critical for separating *R. japonicus* from the morphologically similar *R. crispus* and *R. obtusifolius* . Representative images of these characters were taken.

### Plant Materials and Growth Conditions

Seeds of *R. japonicus, R. crispus, R. obtusifolius* were sown on moist filter paper in Petri dishes and incubated under a 12 h light / 12 h dark photoperiod at 25/15 °C (day/night) until germination. Seedlings were then transplanted into plastic pots ( φ = 5 cm), with one seedling per pot, and maintained under the same environmental conditions. For symptom reproduction assays, a commercial potting soil (Tanemaki Baido; Takii & Co., Ltd., Japan) was used. For quantitative pathogenicity assays, seedlings were grown in a soil mixture of akadama soil and vermiculite (1:1, v/v).

### Inoculum Preparation

Inoculum preparation followed the method described by Izumi and Sato (2025). Briefly, strain GR1 was pre-cultured for 4 days in a 300 mL Erlenmeyer flask containing 50 mL of 0.5% yeast extract–malt (YM) liquid medium by inoculating a 5-mm-diameter mycelial disc, and incubated on a rotary shaker at 25 °C and 130 rpm. The main culture was initiated by transferring 1% (v/v) of the preculture into a 500 mL Sakaguchi flask containing 150 mL of YM liquid medium and incubated for 7 days at 25 °C and 130 rpm. The resulting culture was filtered through two layers of gauze to remove large mycelial fragments. Fungal suspensions were prepared from the filtrate and adjusted using YM liquid medium to a concentration of 1.0 × 10⁵ propagules mL⁻¹ for symptom reproduction assays or 1.0 × 10⁶ propagules mL⁻¹ for quantitative pathogenicity assays. Here, the term “propagules” refers to conidia and small mycelial fragments.

### Pathogenicity Assay on *R. japonicus* for Symptom Reproduction

*R. japonicus* plants grown for approximately one month were sprayed with the fungal suspension (1.0 × 10⁵ propagules mL⁻¹) until runoff. After inoculation, plants were enclosed in transparent polyethylene bags to maintain high humidity and incubated in darkness at 30 °C for 48 h. Plants were then transferred to an acrylic growth box and maintained at 30 °C under a 12 h light/12 h dark photoperiod with 90 ± 10% relative humidity.

Disease development was monitored regularly, and representative symptoms were photographed 10 days after inoculation. Magnified images of representative disease symptoms were obtained using a digital microscope (VHX-970F, Keyence, Japan). To re-isolate the pathogen, conidia produced on symptomatic leaf surfaces were picked up with a sterile inoculating loop and streaked onto PDA plates. Plates were incubated at 25 °C in the dark, and emerging fungal colonies were examined morphologically, including colony and conidial characteristics, to confirm identity.

### Quantitative Pathogenicity Assay on *Rumex* species

Pathogenicity of strain GR1 was evaluated on three *Rumex* species (*R. japonicus*, *R. crispus*, and *R. obtusifolius*) as previously described, with minor modifications (Izumi and Sato 2025). Seedlings at the 1–2 leaf stage were sprayed with a suspension (1.0 × 10⁶ propagules mL⁻¹) at an equivalent spray volume of 4000 L ha⁻¹. Control plants were sprayed with sterile YM medium. After inoculation, plants were placed in humid chambers created using moistened polyethylene bags and incubated at 25 °C in darkness for 48 h. Plants were then moved to standard growth conditions (25 °C, 12 h light/12 h dark), while high humidity was maintained by keeping the polyethylene bags in place. Leaf-level disease incidence was recorded 21 days after inoculation, and shoot biomass of *Rumex* plants was measured as fresh weight at the same time point. Leaf-level disease incidence was expressed for each plant as the proportion of leaves showing clear disease symptoms relative to the total number of leaves. A leaf was scored as diseased when characteristic signs were observed (e.g., visible white conidiomata).

### Statistical Analysis

Statistical analyses were conducted in R (version 4.5.0). One plant grown in one pot was treated as one biological replicate. For each *Rumex* species, *n* = 4–5 plants per treatment (GR1 inoculation vs. mock control) were used, and the bioassay was repeated twice on separate dates. Leaf-level disease incidence was compared between treatments using the Mann–Whitney U test. Shoot biomass was analyzed using Student’s *t*-test after confirming homogeneity of variances with an F-test. Data are presented as mean ± SE. Similar results were obtained in the second run, and data from one representative run are shown.

## Results

### Morphological Characterization of Strain GR1

When cultured on PDA at 25 °C in the dark, GR1 developed very slowly, with colonies attaining only ∼20 mm in diameter after 33 days ( Fig. 1). The colonies were round in outline but displayed an uneven, elevated surface with coarse folds and wrinkles. The upper surface was distinctly black at the center and surrounded by a white marginal zone. A reddish pigment diffused into the agar, particularly around the colony center. On the reverse, the colony center was dark reddish -brown and contrasted with a paler peripheral zone. Microscopically, the fungus produced hyaline, cylindrical conidia that were mostly aseptate, though occasional 1 -septate forms were observed, and had slightly rounded ends. Conidia measured 6.9–21.9 × 1.3–3.1 µm (*n* = 50) (Fig. 2).

**Figure 1.**
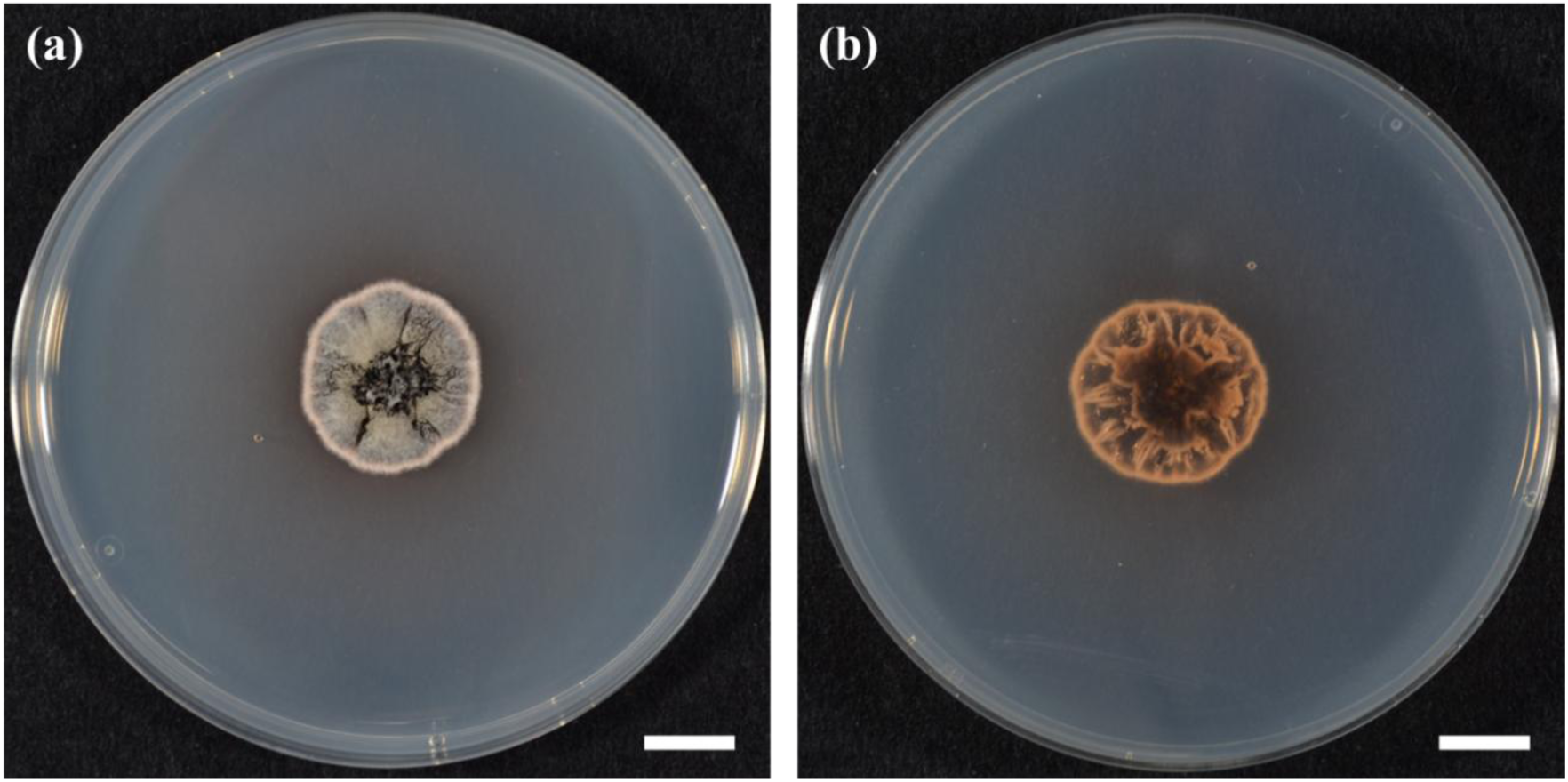
**Colony morphology of strain GR1.** Strain GR1 cultured on potato dextrose agar (PDA) for 1 month at 25 °C in darkness. (a) Colony surface; (b) colony reverse. Scale bars: 10 mm.

**Figure 2.**
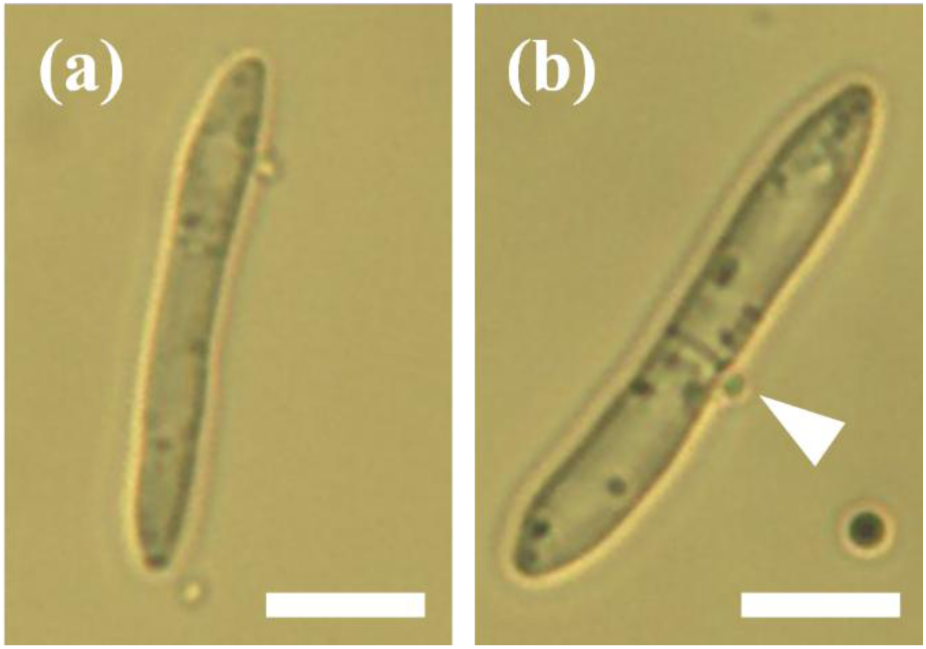
**Conidial morphology of strain GR1.** Light micrographs showing conidia produced by strain GR1: (a) a typical aseptate conidium; (b) a rare 1-septate conidium (arrowhead). Conidia were cylindrical and hyaline. Scale bars = 5 µm.

### Molecular Identification of Strain GR1

Comparison of the ITS sequence of GR1 with the NCBI database revealed identical to several *T. rumicicola* entries (PX069134, OL711641, PP301328, PP301329) as well as to the ex-type of *T. rumicis* (NR_174642). The *T. rumicicola* ex-type strain (NR_154517) also showed nearly identical similarity (469 of 471 bp; 99.6%). For the LSU locus, GR1 exhibited 100% identity to the *T. rumicicola* ex-type strain and three additional isolates (KX287254–KX287256, PX069135), whereas the match to *T. rumicis* was 98.6% (697/707 bp; MW276098). Two further *T. rumicicola* accessions (PP301336, PP301337) were also highly similar (99.9%, 722/723 bp).

A maximum-likelihood phylogeny constructed from a concatenated ITS-LSU alignment (1,191 bp) showed four major clades within *Teratoramularia*: *T. kirschneriana*, *T. infinita*, *T. persicariae*, and a mixed *T. rumicicola*-*T. rumicis* clade (Fig. 3). Within this latter group, GR1 consistently grouped with *T. rumicicola* isolates and the *T. rumicicola* ex-type, with full bootstrap support (100%). Although the *T. rumicis* type strain was retained in the same clade, it was placed on a noticeably extended branch, reflecting divergence from GR1 and the *T. rumicicola* assemblage.

**Figure 3.**
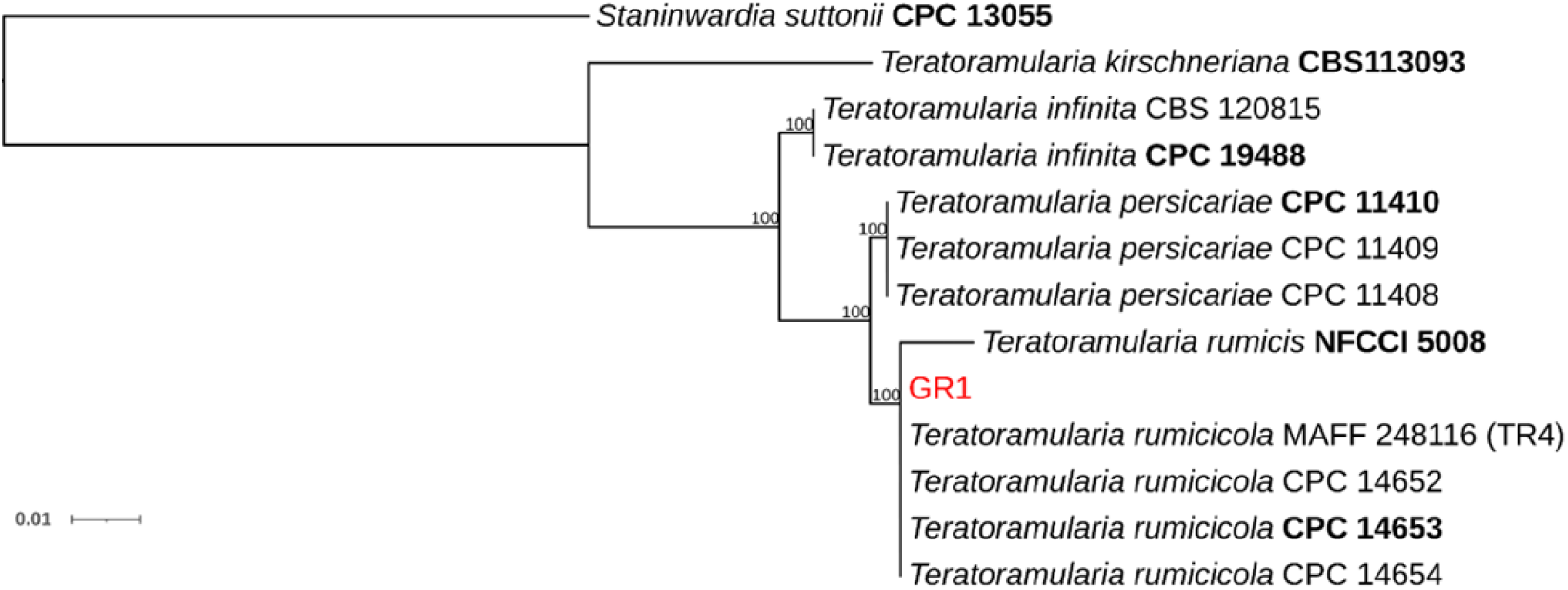
**Phylogenetic position of strain GR1 inferred from concatenated ITS and LSU sequences.** Maximum-likelihood phylogeny reconstructed from concatenated ITS–LSU rDNA sequences under the GTR+GAMMA substitution model. Node labels indicate bootstrap support (1,000 replicates), with values ≥70% shown. GR1 is highlighted in red; ex-type strains are indicated in bold. Reference sequences of *Teratoramularia* spp. were obtained from GenBank. *Staninwardia suttonii* served as the outgroup.

### Identification of Host Plant Voucher Specimen

The overall morphology of the inflorescence and leaves is shown in Fig. 4a. The leaf base is cuneate, lacking auricles on both sides, thus indicating subgenus *Rumex* (Fig. 4a). As shown in Fig. 4b, the margin of the inner tepal is shallowly serrate, a key diagnostic character for distinguishing *R. japonicus* from *R. crispus*, which has entire margins, and *R. obtusifolius*, which has sharply serrate margins. Taken together, these diagnostic characters support the identification of the voucher specimen as *R. japonicus*. Notably, whitish fungal residue was visible on the surface of the specimen (Fig. 4c), which is consistent with fungal growth observed in *T. rumicicola* infections.

**Figure 4.**
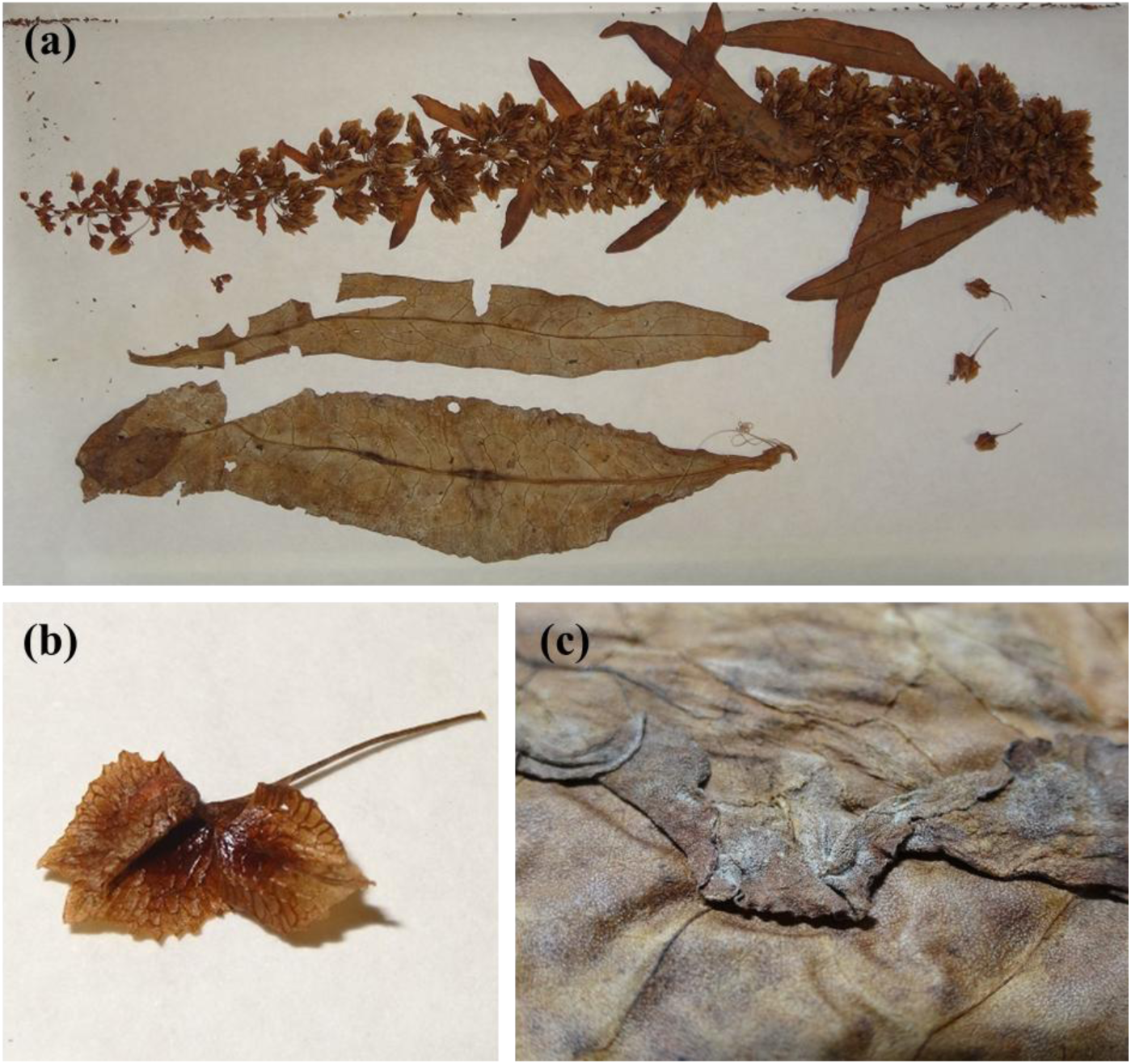
**Host plant voucher specimen from which strain GR1 was isolated.** (a) Overall specimen showing the inflorescence and leaves. (b) Inner tepal showing a shallowly serrate margin, diagnostic of *Rumex japonicus*. (c) White fungal residue on the leaf surface.

### Symptom reproduction on *R. japonicus*

A photograph of the original field symptom, taken by coauthor T. Sato in 1996 and archived in the NARO Genebank, showed leaves extensively covered with conspicuous white, powdery fungal growth (Fig. 5a; reprinted from the Microorganisms image database, NARO Genebank, Japan). In the present inoculation assay, similar symptoms developed on *R. japonicus*, and marked white fungal growth was evident on leaves by 10 days after inoculation (Fig. 5b and 5c). Accordingly, the inoculation assay reproduced the original symptom. To confirm the causal relationship, conidia were collected directly from symptomatic leaf surfaces and cultured on PDA. The resulting colonies and conidial morphology were indistinguishable from those of the original isolate GR1, supporting that GR1 was the causal agent of the symptoms observed on *R. japonicus*.

**Figure 5.**
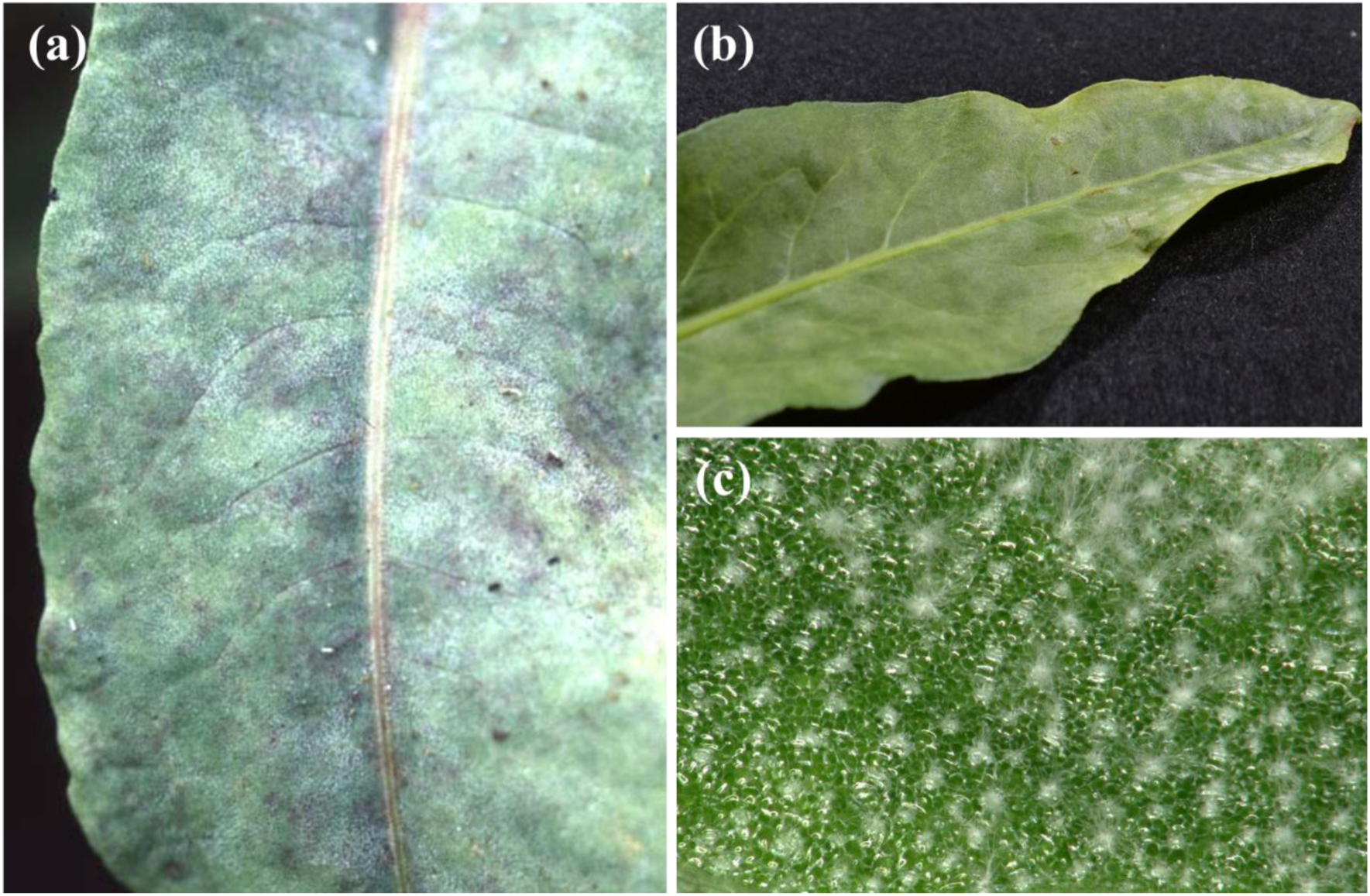
**Comparison of natural and reproduced symptoms of GR1 on *R. japonicus*.** (a) Naturally infected leaf collected in Hahajima, the Bonin Islands (1996), showing white powdery symptoms on the leaf surface (original symptoms; source of strain GR1). The photograph is reprinted from the Microorganisms image database, NARO Genebank, Japan (https://www.gene.affrc.go.jp/databases-micro_images_detail.php?id=25206 ). (b) Leaf 21 days after inoculation with GR1 (this study), showing white powdery symptoms consistent with the natural symptoms. (c) Higher-magnification view of the leaf surface showing the white powdery mold masses.

### Quantitative pathogenicity and host range within three *Rumex* species

Twenty one days after inoculation, leaf-level disease incidence reached 84.0 ± 4.0% in *R. crispus* and 84.0 ± 4.0% in *R. japonicus*, whereas it was lower in *R. obtusifolius* (40.0 ± 6.7%) (Fig. 6). Consistent with symptom development, shoot fresh weight was reduced in inoculated plants across all three species, with significant growth inhibition of 44.2 ± 5.0% in *R. crispus*, 56.1 ± 7.3% in *R. japonicus*, and 28.7 ± 9.2% in *R. obtusifolius*. Overall, *R. obtusifolius* showed comparatively lower susceptibility to GR1.

**Figure 6.**
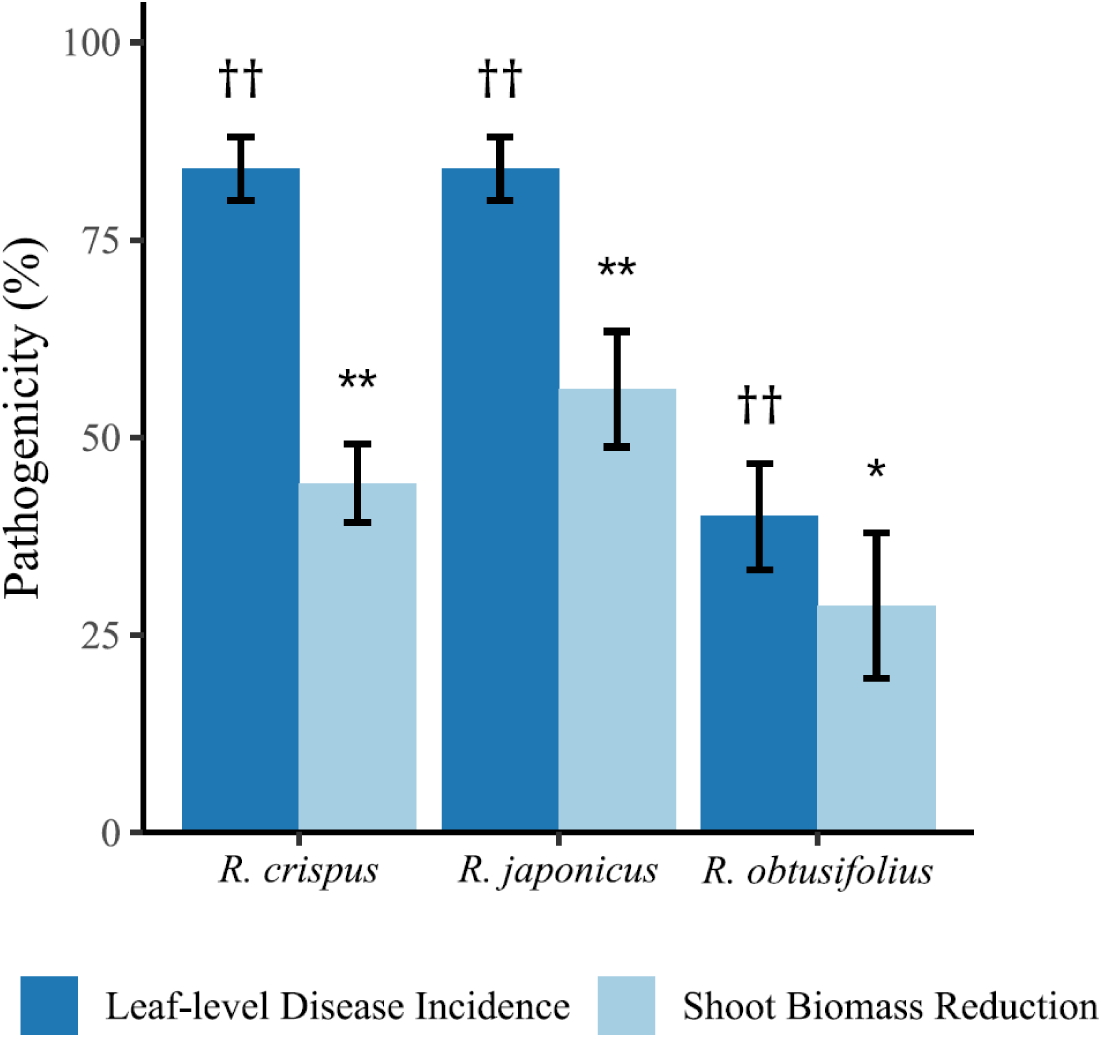
**Pathogenicity of strain GR1 on three *Rumex* species 21 days after inoculation.** Bars show mean ± SE ( *n* = 4-5). Dark blue: leaf-level disease incidence; light blue: shoot biomass reduction relative to the mock control. Daggers (††, *p* < 0.01) and asterisks (* *p* < 0.05; ** *p* < 0.01) indicate significant differences versus the corresponding controls within each species (incidence: Mann–Whitney U test, biomass: Student’s *t*-test after an F-test for homoscedasticity).

## Discussion

Strain GR1 was identified as *T. rumicicola* based on concordant morphological, molecular, phylogenetic, and biogeographic evidence. The colony phenotype of GR1 and the shape of its conidia closely matched the diagnostic features reported for the *T. rumicicola* ex-type strain (CPC 14653) (Videira et al. 2016). In contrast, the closest related species, *T. rumicis*, is typically characterized by longer conidia and a higher frequency of septate conidia (0 –2 septa) (Verma et al. 2021), which were not predominant in GR1. Molecular data provided additional support. The ITS and LSU sequences of GR1 was identical or nearly identical to that of the *T. rumicicola* ex-type strain. In phylogenetic analysis of concatenated ITS–LSU datasets, GR1 consistently clustered with reference isolates of *T. rumicicola*, including the ex-type, and was diverged from *T. rumicis*. Moreover, *T. rumicicola* has been documented repeatedly from East Asia (including Korea and Japan), whereas *T. rumicis* has so far been reported only from a single collection in India (Videira et al. 2016; Verma et al. 2021; Choi et al. 2025; Izumi and Sato 2025). Because multi-locus sequences for *T. rumicis* (e.g., RPB2, TEF1) are not currently available, species boundaries within this complex cannot be evaluated as rigorously as would be possible with additional loci. Nonetheless, the congruence of morphological traits, ITS –LSU similarity, phylogenetic position, and presently known biogeography provides a robust support for identifying GR1 as *T. rumicicola*.

The re-identification of strain GR1, isolated in 1996, indicates that *T. rumicicola* was already present in Japan approximately two decades before its formal description. The isolate underpinning the original description was obtained in 2007 (Videira et al. 2016), and subsequent records include isolates collected in 2022 (Choi et al. 2025) and 2024 (Izumi and Sato 2025). Thus, to our knowledge, GR1 represents the earliest documented record of this species. Notably, GR1 originated from Hahajima in the Bonin Islands, a remote oceanic archipelago geographically isolated from the Japanese mainland (Fig. 7). While an introduction from outside the islands cannot be ruled out, the presence of GR1 in 1996 raises the possibility that *T. rumicicola* had persisted in the local ecosystem for an extended period. In addition, GR1 represents a first record from a subtropical island environment. To our knowledge, previous isolates of *T. rumicicola* have been reported from temperate regions, and therefore this finding extends the known ecological and geographic context of the species. Accumulating such records across climatic zones will be important for refining our understanding of the distribution of *T. rumicicola*. Because GR1 originated from an isolated subtropical oceanic archipelago, targeted surveys across subtropical and temperate islands and mainland sites will be needed to evaluate the broader distribution of *T. rumicicola*.

**Figure 7.**
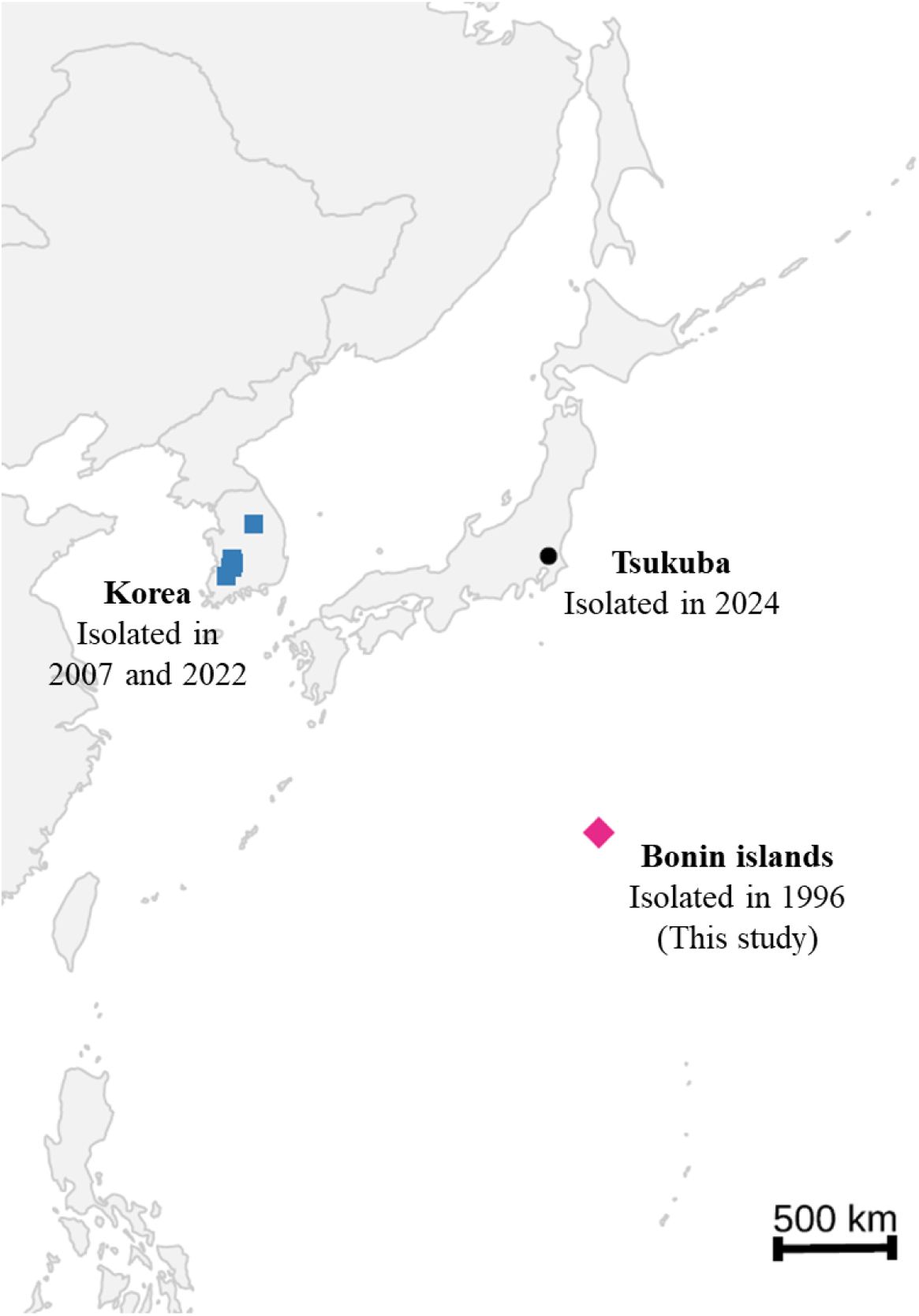
**Geographic distribution of *T. rumicicola* isolates in East Asia.** The black circle indicates Tsukuba, Japan (isolated in 2024; Izumi and Sato, 2025). The magenta diamond indicates the Bonin Islands (isolated in 1996; this study). Blue squares indicate Korean localities (isolated in 2007 and 2022 ; Videira et al., 2016; Choi et al., 2025). A 500 km scale bar is shown. Base map: U.S. Census Bureau / *maps* package (public domain).

In addition to the morphological identification of the host plant as *R. japonicus*, we examined historical distribution records to assess whether this species could plausibly have been the natural host at the time and location of isolation. Herbarium records document the presence of *R. japonicus* on Hahajima as early as 1920 (e.g., specimen TI:ISP:576; GBIF Japan), well before the 1996 isolation of strain GR1. The species was also included in the revised flora of the Bonin and Volcano Islands (Kobayashi and Ono 1987), supporting its long -established occurrence within the archipelago. In contrast, *R. crispus* has been documented from the Bonin Islands since 2005 from Chichijima (Ogasawara World Natural Heritage Center 2013). In particular, *R. japonicus* is recognized as a native species in Japan, whereas *R. crispus* is regarded as an alien species; thus, the early establishment of *R. crispus* in the remote Bonin Islands is considered unlikely. Taken together, these lines of evidence support the conclusion that the natural host of strain GR1 was *R. japonicus*. Importantly, the present study establishes *R. japonicus* as the first confirmed natural host of *T. rumicicola*. Clarifying the natural host range of *T. rumicicola* will be essential for understanding its behavior within native and managed ecosystems. Finally, based on the characteristic white, powdery growth on leaf surfaces, we propose the name white powdery blight (Japanese: hakufun -byo) for this disease.

Beyond its natural host, GR1 also showed pathogenicity toward other *Rumex* weeds, including *R. crispus* and *R. obtusifolius*, indicating that susceptible hosts extend beyond *R. japonicus* under experimental conditions. This pattern is consistent with our previous report on a different Japanese isolate of *T. rumicicola*. Although further work is required to evaluate susceptibility of other *Rumex* species and related genera within Polygonaceae, these findings support the potential utility of *T. rumicicola* as candidates for *Rumex* weed management.

We re-evaluated the archival strain GR1 (Bonin Islands, 1996; NARO Genebank) and identified it as *T. rumicicola*. This represents the earliest documented record of the species, confirms *R. japonicus* as its first verified natural host, and extends its known distribution to a subtropical region.

## Author Contributions

M.I. conceived and designed the study, performed the experiments, analyzed the data, and drafted the manuscript. T.S. contributed to taxonomic assessment, provided resources (archived strain and symptom photographs deposited in the NARO Genebank), and advised on the experimental conditions for reproducing original symptoms. Both authors reviewed and edited the manuscript and approved the final version.

## Acknowledgements

This work was partially supported by JSPS KAKENHI Grant Numbers 24KJ1012 , which granted to M. I. We sincerely thank Dr. Mitsuo Horita, the curator of the plant pathological herbarium, the Institute for Agro-Environmental Sciences, NARO, Japan for lending the voucher specimen.

## Disclosure statement

The authors declare no conflict of interests.

## Notes

### Competing Interest Statement

The authors have declared no competing interest.

## References

1. Bond, W., Davies, G., & Turner, R. J. 2007. “The Biology and Non -Chemical Control of Broad-Leaved Dock (Rumex obtusifolius L.) and Curled Dock (R. crispus L.).” HDRA, Ryton Organic Gardens, UK. 2007. https://garden-organic.files.svdcdn.com/production/documents/dock-review.pdf?dm=1726651587.

2. Choi, Joon-Ho, Young-Joon Choi, Lamiya Abasova, Hye-Ryeong Jang, Hyeon-Dong Shin, and Ji-Hyun Park. 2025. “ First Report of a Powdery Mildew - like Disease Caused by *Teratoramularia rumicicola* on *Rumex obtusifolius* in Korea.” Journal of Phytopathology 173 (3). 10.1111/jph.70103.

3. Izumi, M., and Sato, T. 2025. “Bioherbicidal Activity and Host Range of *Teratoramularia rumicicola* Strain TR4 Isolated From *Rumex crispus* in Japan.” Weed Biology and Management 25 (4): e70008.

4. Letunic, Ivica, and P. Bork. 2024. “Interactive Tree of Life (ITOL) v6: Recent Updates to the Phylogenetic Tree Display and Annotation Tool.” Nucleic Acids Research 52 (April): W78–82.

5. McGrann, Graham R. D., Ambrose Andongabo, Elisabet Sjökvist, Urmi Trivedi, Francois Dussart, Maciej Kaczmarek, Ashleigh Mackenzie, et al. 2016. “The Genome of the Emerging Barley Pathogen Ramularia Collo-Cygni.” BMC Genomics 17 (1): 584.

6. Ogasawara World Natural Heritage Center. 2013. “Issue Analysis and Reference Materials for Developing an Action Plan to Prevent Invasion and Spread of New Alien Species in the Ogasawara Islands [in Japanese].” https://ogasawara-info.jp/wp-content/uploads/2024/10/3-2_Issue_Analysis_and_Reference_Materials_for_Developing_Action_Plan_to_Prevent_Invasion_and_Spread_of_New_Alien_Species_2013.pdf

7. Ohashi, H., Kadota, Y., Murata, J., Yonekura, K., & Kihara, H. (eds.)., ed. 2017. Wild Flowers of Japan, Vol. 4: Malvaceae–Apocynaceae. Tokyo, Japan: Heibonsha.

8. Sato, T., Uzuhashi, S., Hosoya, T., and Hosaka K. 2010. “A List of Fungi Found in the Bonin (Ogasawara) Islands.” Ogasawara Research, no. 35 (March): 59–160.

9. Stamatakis, A. 2014. “RAxML Version 8: A Tool for Phylogenetic Analysis and Post - Analysis of Large Phylogenies.” Bioinformatics 30 (January): 1312–13.

10. Tamura, K., G. Stecher, and Sudhir Kumar. 2021. “MEGA11: Molecular Evolutionary Genetics Analysis Version 11.” Molecular Biology and Evolution 38 (April): 3022–27.

11. Verma, S. K., P. Kushwaha, S. K. Yadav, and Raghvendra Singh. 2021. “Morphology and Phylogeny of *Teratoramularia rumicis* —a New Foliar Pathogen of *Rumex crispus* from India and Diversity of Ramularioid Complex on Rumex Spp.” Phytotaxa 523 (October): 208–28.

12. Videira, S., J. Z. Groenewald, U. Braun, Hyeon - dong Shin, P. W. Crous, and P. W. Crous. 2016. “All That Glitters Is Not Ramularia.” Studies in Mycology 83 (March): 49–163.

13. Vilgalys, R., and M. Hester. 1990. “Rapid Genetic Identification and Mapping of Enzymatically Amplified Ribosomal DNA from Several Cryptococcus Species.” Journal of Bacteriology 172 (8): 4238–46.

14. White, T. J., T. Bruns, Lee S J W, and J. Taylor. 1990. “Amplification and Direct Sequencing of Fungal Ribosomal RNA Genes for Phylogenetics.” PCR Protocols: A Guide to Methods and Applications 18 (1): 315–22.

