## Supplemental Table 1 for "Early occurrence of *Teratoramularia rumicicola* in Japan: re-identification of historic strain GR1 and its pathogenic effects on *Rumex* species"

**Supplementary Table 1. GenBank accessions for ITS and LSU rDNA sequences included in the phylogenetic analyses.**

The table summarizes ITS and LSU sequences retrieved from GenBank, together with strain information and source references. Sequences generated and analyzed in the present study are highlighted in bold.

| Species | Culture number | ITS | LSU | References |
|---|---|---|---|---|
| *Teratoramularia infinita* | CBS 120815 | KX287544 | KX287248 | Videira et al. 2016 |
| | CBS 141104; CPC 19488 | KX287545 | KX287249 | Videira et al. 2016 |
| *Teratoramularia kirschneriana* | CBS 113093; RoKi 1144 | GU214669 | GQ852627 | Crous et al. 2009 |
| *Teratoramularia persicariae* | CPC 11408 | KX287546 | KX287250 | Videira et al. 2016 |
| | CPC 11409 | KX287547 | KX287251 | Videira et al. 2016 |
| | CBS 141105; CPC 11410 | KX287548 | KX287252 | Videira et al. 2016 |
| *Teratoramularia rumicis* | NFCCI 5008 | MK967558 | MW276098 | Verma et al. 2021 |
| ***Teratoramularia rumicicola*** | **GR1** | **PX601562** | **PX599000** | **This study** |
| | MAFF 248116 (TR4) | PX069134 | PX069135 | Izumi & Sato 2025 |
| | CPC 14652 | KX287549 | KX287254 | Videira et al. 2016 |
| | CBS 141106; CPC 14653 | KX287550 | KX287255 | Videira et al. 2016 |
| | CPC 14654 | KX287551 | KX287256 | Videira et al. 2016 |
| *Staninwardia suttonii* | CPC 13055 | DQ923535 | DQ923535 | Summerell et al. 2006 |

### References


Crous, P. W., C. L. Schoch, K. D. Hyde, A. R. Wood, C. Gueidan, G. S. de Hoog, and J. Z. Groenewald. 2009. "Phylogenetic Lineages in the Capnodiales." *Studies in Mycology* 64 (June): 17-47S7.

Izumi, M., and Sato, T. 2025. "Bioherbicidal Activity and Host Range of *Teratoramularia rumicicola* Strain TR4 Isolated From *Rumex crispus* in Japan." *Weed Biology and Management* 25 (4): e70008.

Summerell, B., J. Groenewald, A. Carnegie, R. Summerbell, and P. Crous. 2006. "Eucalyptus Microfungi Known from Culture. 2. Alysidiella, Fusculina and Phlogicylindrium Genera Nova, with Notes on Some Other Poorly Known Taxa." *Fungal Diversity* 23: 323–50.

Verma, S. K., P. Kushwaha, S. K. Yadav, and Raghvendra Singh. 2021. "Morphology and Phylogeny of *Teratoramularia rumicis* —a New Foliar Pathogen of *Rumex crispus* from India and Diversity of Ramularioid Complex on *Rumex* Spp." *Phytotaxa* 523 (October): 208–28.

Videira, S., J. Z. Groenewald, U. Braun, Hyeon-dong Shin, P. W. Crous, and P. W. Crous. 2016. "All That Glitters Is Not *Ramularia*." *Studies in Mycology* 83 (March): 49–163.
